# Visualizing H_2_O_2_ and NO in endothelial cells: strategies and pitfalls

**DOI:** 10.1101/2023.02.15.528776

**Authors:** Hamza Yusuf Altun, Melike Secilmis, Tuba Akgul Caglar, Emre Vatandaslar, Gürkan Öztürk, Sven Vilain, Emrah Eroglu

## Abstract

The relationship between hydrogen peroxide (H_2_O_2_) and nitric oxide (NO) in the vasculature is multifaceted and remains controversial because the dynamic detection of these reactive molecules is challenging. Genetically encoded biosensors (GEBs) allow visualizing real-time dynamics in living cells and permit multiparametric detection of several analytes. Although robust, GEBs’ utility depends on several parameters that need fine-tuning for proper imaging and correct data analysis: i.e., camera binning, temperature, and the resolution power of the imaging instruments are some critical parameters that require optimization. We have generated a new double-stable transgenic endothelial cell line stably expressing the biosensors HyPer7 and O-geNOp and systematically tested different imaging modes and their impact on the performance of each biosensor. Ambient temperature and the type of imaging mode did not influence the results, while camera resolution settings significantly affected readouts of HyPer probes but not O-geNOp. Changing a single parameter in a co-imaging mode significantly altered the biosensor’s dynamic measurements, potentially causing misinterpretation. This study provides a general guide and the pitfalls of employing GEBs in a multispectral imaging mode.

## 1. Introduction

The relatively stable reactive oxygen species (ROS), hydrogen peroxide (H_2_O_2_), and the short-lived radical nitric oxide (NO) are both signaling molecules that play essential roles in the function of vascular cells [1]–[3]. Various cells produce H_2_O_2_ as a byproduct of normal cellular metabolism [4]. In response to various stimuli, such as inflammation or oxidative stress [5]–[7], H_2_O_2_ can act as a signaling molecule and can activate certain enzymes and regulate gene expression [8]. A group of specialized and tissue-specific isozymes, NO synthases (NOS), can dynamically generate NO [9]. In the vascular systems, the endothelial isoform NOS (eNOS) regulates blood pressure and relaxes smooth muscle cells [10]–[12].

The relationship between H_2_O_2_ and NO in vascular cells is complex and multifaceted [13]–[17] and experimentally tricky to analyze due to the lack of established techniques. In this regard, genetically encoded biosensors (GEBs) are the preferred technique among potential tools for dynamically monitoring the interaction between H_2_O_2_ and NO in living cells [18], [19]. One of the most significant advantages of GEBs is that they allow for studying dynamic processes in living cells [20]. These probes provide a high level of specificity and sensitivity compared to other methods and are minimally invasive [21]. Because GEBs are synthesized by cells as proteins, they are less intrusive than other methods, which makes them helpful in studying biological processes in living cells over long periods [22]. Also, GEBs can be readily introduced into cells using various methods, including transfection, viral infection, and gene editing [23].

Among various biosensors, we selected the green fluorescent protein (GFP)-based H_2_O_2_ sensitive biosensor HyPer7 [24] and the orange fluorescent protein (OFP)-based NO probe O-geNOp [25]–[27] to simultaneously measure these two reactive molecules in vascular endothelial cells using multispectral imaging approaches. However, endothelial cells are known to be difficult to transfect [28]. To overcome these challenges, we have recently developed new strategies to improve transfection efficiency in endothelial cells by using viral gene delivery methods [23], [28].

Another important fact in employing GEBs in live-cell imaging is the requirement of several optimization steps [29]. One key factor affecting the performance of GEBs is the signal-to-noise ratio (SNR). Camera binning is one parameter that can be tuned in live-cell imaging to improve the sensitivity and SNR [30]. It combines the charge from multiple pixels into a single “binned” pixel, resulting in a higher SNR and a lower background noise level [30]. Camera binning increases the SNR and requires lower exposure times to capture an image. Illuminating the specimens with lower power can be an advantage for live-cell imaging, as longer exposure times may increase the likelihood of photobleaching or phototoxicity, which often comes with ROS generation [31], [32]. Thus, camera binning can be a valuable tool for improving fluorescence microscopy’s sensitivity and SNR. Overall, the imaging system’s resolution is an essential factor that can affect the performance of GEBs, which is critical in choosing imaging instruments such as epifluorescence-, spinning disk-, or confocal microscopes.

Temperature is another critical factor in live-cell imaging that affects the model systems’ functionality and can significantly affect the performance of GEBs. During live-cell experiments, mammalian cells should ideally be kept at 37°C, which may not be optimal for the functionality of specific GEBs [33], [34]. Thus, it is essential to consider the role of temperature and optimize the conditions for each experiment.

Accurate detection of ROS and RNS dynamics can be critical in understanding the relationship between H_2_O_2_ and NO under normal and pathological conditions. This study generated a new endothelial cell line stably co-expressing HyPer7 and O-geNOp. We performed live-cell imaging experiments to optimize and understand the dynamics of HyPer7 and O-geNOp with different camera binning setups and at various temperatures using special imaging rigs with different resolution capacities. Overall, this study guides how to reliably co-image intracellular H_2_O_2_ and NO production.

## 2. Results

### 2.1. Generation of a double-stable endothelial cell line expressing HyPer7 and O-geNOp

To simultaneously assess cellular levels of H_2_O_2_ and NO in endothelial cells, we generated a transgenic cell line expressing the biosensors HyPer7 and O-geNOp using lentivirus transduction in immortalized endothelial cells EA.hy926 (Figure 1A). Using fluorescence-assisted cell sorting (FACS), we selected a cell population expressing both biosensors and detected robust expression levels in 100% of the cells (Figure 1B). Next, we tested the functionality of the biosensors in this double-stable cell line by administering low levels of exogenous H_2_O_2_ or NO by imaging the biosensors individually. Due to the ratiometric nature of HyPer7, H_2_O_2_ provision caused an instant decrease in the blue excitation channel (Ex: F420 and Em: F525) and a simultaneous increase in the green excitation channel (Ex: F475 and Em: F525), confirming the functionality of the biosensor and permitting ratiometric imaging of intracellular H_2_O_2_ levels (Figure 1C). The lower panel of Figure 1C shows HyPer7’s operation, which consists of a circularly permuted green fluorescent protein (cpGFP) and H_2_O_2_-sensitive subunits termed OxyR. Next, we examined the functionality of O-geNOp by providing the NO-amine complex NOC-7 [40]. Besides HyPer7, as shown in the lower panel of Figure 1D, geNOp biosensors are single FP-based probes that are intensiometric and reversible. As expected, the O-geNOp signal (Ex: F555 and Em: F605) instantly changed upon administration of the NO-releasing compound (Figure 1D). This approach showed that hardly transfectable endothelial cell lines are robustly transducable to yield functional biosensors allowing real-time detection of H_2_O_2_ and NO.

**Figure 1:**
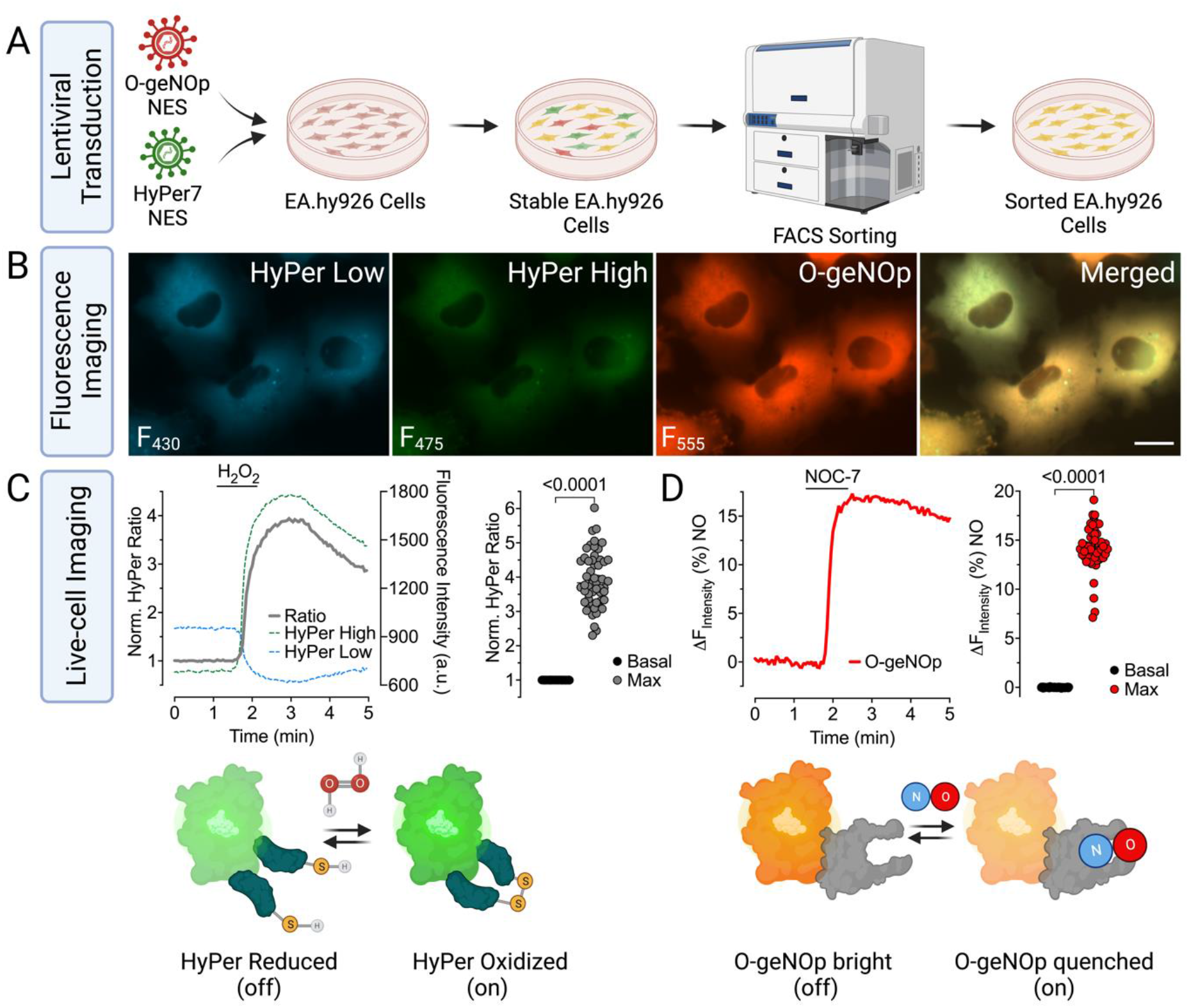
Schematic overview and workflow for developing endothelial cells stably co-expressing HyPer7 and O-geNOp **(A)** Native endothelial cells (EA.hy926) at low passage were transduced with lentivirus containing ORFs of HyPer7 and O-geNOp, respectively. 48-72h after transduction, HyPer7 and O-geNOp expressing cells were sorted using FACS. **(B)** Representative widefield images of endothelial cells after FACS. The first three panels show images of cells under different optical setups according to the optical properties of the biosensors HyPer Low (Ex/Em: 430 nm/525 nm) (first image), HyPer High (Ex/Em: 475 nm/525 nm) (second image), O-geNOp (Ex/Em: 555 nm/605 nm) (third image) and fourth image shows merged channels. The scale bar represents 20 μm. **(C)** The representative curve shows HyPer7 signals upon administration of 50 μM H_2_O_2_. The scatter dot plot shows basal levels (black dots, n=3/57) and maximum HyPer7 responses (grey dots, n=3/57) of individual cells. The lower scheme shows the working principle of the HyPer7 biosensor. **(D)** Representative O-geNOp signals show cell responses after the provision of 10 μM NOC-7. The scatter dot plot shows O-geNOp responses under basal levels (black dots, n=3/57) and maximum responses upon administration of 10 μM NOC-7 (red dots, n=3/57). The lower panel shows the working principle of O-geNOp. (*p-value* < 0.0001, Student’s t-test)

### 2.2. The effect of camera binning settings on HyPer7 and O-geNOp profile

We tried different camera binning configurations to optimize the dynamic responses of HyPer7 and O-geNOp and tested four different camera configurations; 1×1 (no binning), 2×2, 3×3, and 4×4, binning of four, nine, and sixteen adjacent pixels, respectively. First, we assessed the binning effect on HyPer7. This approach showed a decrease in fluorescent image resolution by expanding the number of binned pixels (Figure 2A). Conversely, increasing the binned pixels resulted in a higher H_2_O_2_-induced HyPer7 response (Figure 2B). Unexpectedly, the HyPer7 profiles recovered over a short time to baseline when cells were imaged with a camera binning 1×1. Such a reduction was not observed when more pixels were binned (Figure 2B). Since the signal of HyPer7 is measured as a ratio of high intensity over low intensity, we investigated which signal was mainly affected (i.e., bleaching) by camera binning. While pixel binning did not affect the high-intensity measurements, low-intensity signals were visibly affected by the number of binned pixels (Figure S1). To further study the effect of pixel binning on HyPer7 kinetics, we assessed changes in HyPer7 signals directly after adding exogenous H_2_O_2._ We observed that increasing the binned pixels resulted in a faster measurable response to H_2_O_2_ (Figure 2C). Hence, determining HyPer7 kinetics is dependent on the camera binning configuration. A quantitative assessment of the correlation between the maximum H_2_O_2_-induced HyPer7 signal and the rate of this signal showed that the highest maximum signal ratio and the fastest signal increase were observed with the 4×4 camera binning configuration and the lowest with 1×1 (Figure 2D).

**Figure 2:**
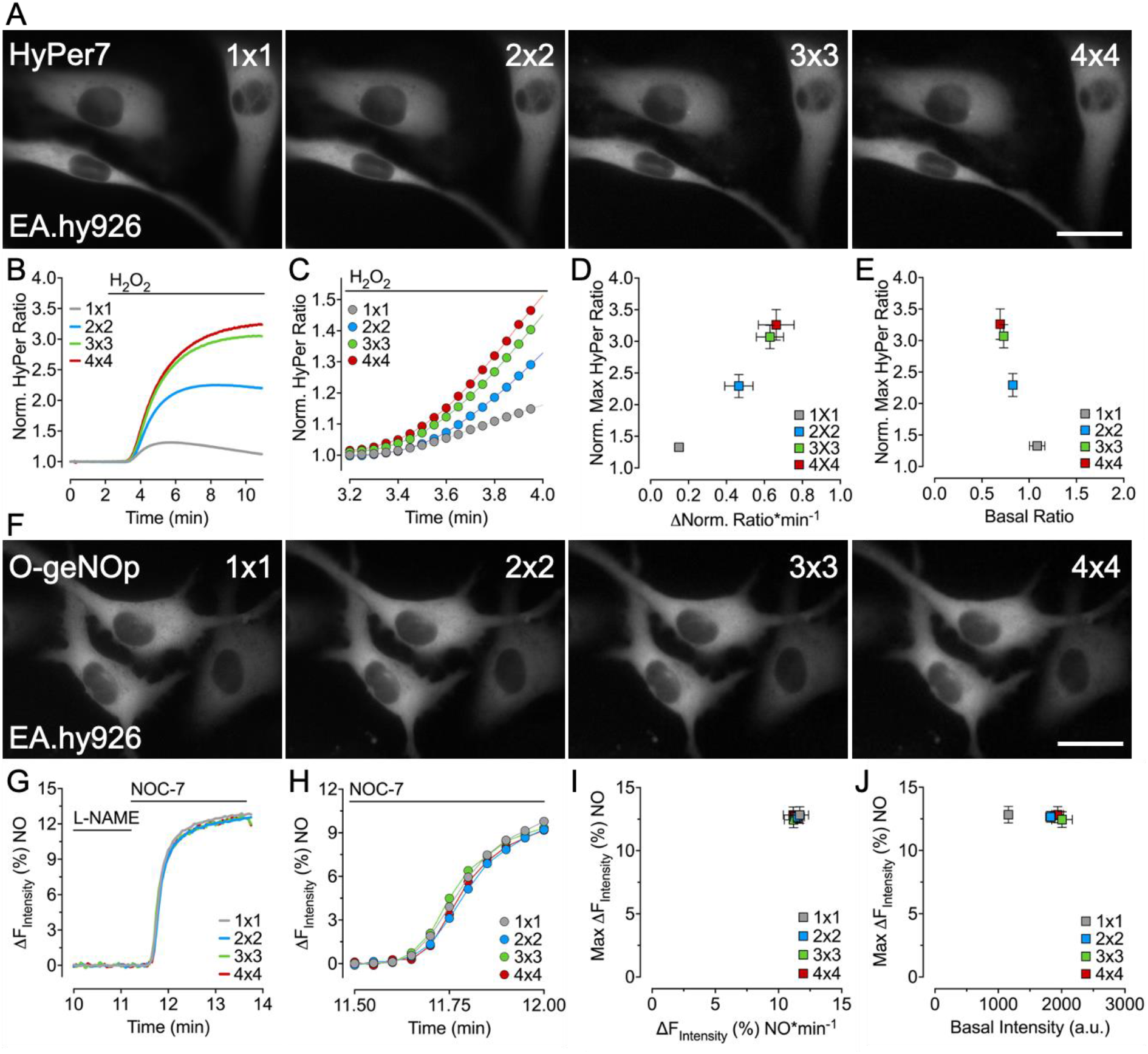
Effect of camera binning settings on HyPer7 and O-geNOp profiles. **(A)** Representative widefield images of EA.hy926 cells stably expressing HyPer7 in the GFP channel. Images were recorded using different camera binning settings 1×1, 2×2, 3×3, and 4×4, as indicated. **(B)** Average curves represent normalized real-time HyPer7 signals in response to 25 μM H_2_O_2_ under different camera binning settings. **(C)** Close-up curves show the dynamics of HyPer7 signals. **(D)** In the XY scatter plot, the Y axis shows the maxima of normalized HyPer7 ratios, and the X axis shows the change of the HyPer7 ratio over time under different camera binning settings. 1×1 (n=6/75), 2×2 (n=6/75), 3×3 (n=6/75), 4×4 (n=6/75). **(E)** In the XY scatter plot, the Y-axis represents the maxima of normalized HyPer ratios. The X-axis shows basal HyPer ratios under different camera binning settings (1×1 (n=6/75), 2×2 (n=6/75), 3×3 (n=6/75), 4×4 (n=6/75), respectively). **(F)** Representative images of cells expressing O-geNOp in different binning configurations **(G)** O-geNOp signals in response to 1 μM of NOC-7 using different camera binning **(H)** Close-up curves show the dynamic of O-geNOp signals **(I)** XY graph was plotted using the rates of O-geNOp fluorescence intensity change with respect to time (X-axis) and maxima of O-geNOp responses (Y-axis) under different camera binning settings. (1×1 (n=6/52), 2×2 (n=6/56), 3×3 (n=6/50), 4×4 (n=6/56). **(J)** XY scatter plot represents the maximum responses of O-geNOp (Y-axis) and basal fluorescent intensity levels (Y-axis). (1×1 (n=6/52), 2×2 (n=6/56), 3×3 (n=6/50), 4×4 (n=6/56). All values are presented as mean ±SEM. Common color code was used for each graphics(Grey (1×1), blue(2×2), green (3×3), red(4×4)). The scale bars represent 20 μm.

Further examination of the relationship between basal HyPer7 ratios and maximum responses showed that the initial basal levels of the HyPer7 signal were close to each other under different camera binning setups (Figure 2E). However, the maximal signal ratio depended on the binning settings (Figure 2E). These observations document that the camera binning setting significantly affects the H_2_O_2_-induced HyPer7 readouts in endothelial cells.

Repeating the same experimental approach in the O-geNOp channel showed decreased image resolution with increased pixel binning, as expected (Figure 2F). Yet, the maximum response to NOC-7 treatment demonstrated O-geNOp’s profile and kinetic was camera binning independent (Figure 2G and H). Accordingly, no significant difference was observed in the signal increase rate and maximum amplitude at different camera binning settings (Figure 2I). Also, there was no observable relationship between the basal levels of O-geNOp and the maximum amplitude (Figure 2J). These experiments show that camera binning does not affect NOC-7-induced O-geNOp signal measurements.

### 2.3. The biosensor’s performance is independent of the resolution power of an imaging rig

Optimal live-cell imaging requires the fast acquisition of images with a high SNR and the highest possible spatial resolution. We tested the biosensor’s performance on conventional widefield (WF) - and spinning disk (SD) microscopes. Fluorescent images in the HyPer7 or O-geNOp channels showed higher contrast and resolution in the SD mode than in WF images (Figures 3A and D). Stimulating HyPer7 with H_2_O_2_ led to a robust increase in the HyPer7 signal with similar profiles and kinetics in both microscopy settings (Figure 3B).

**Figure 3:**
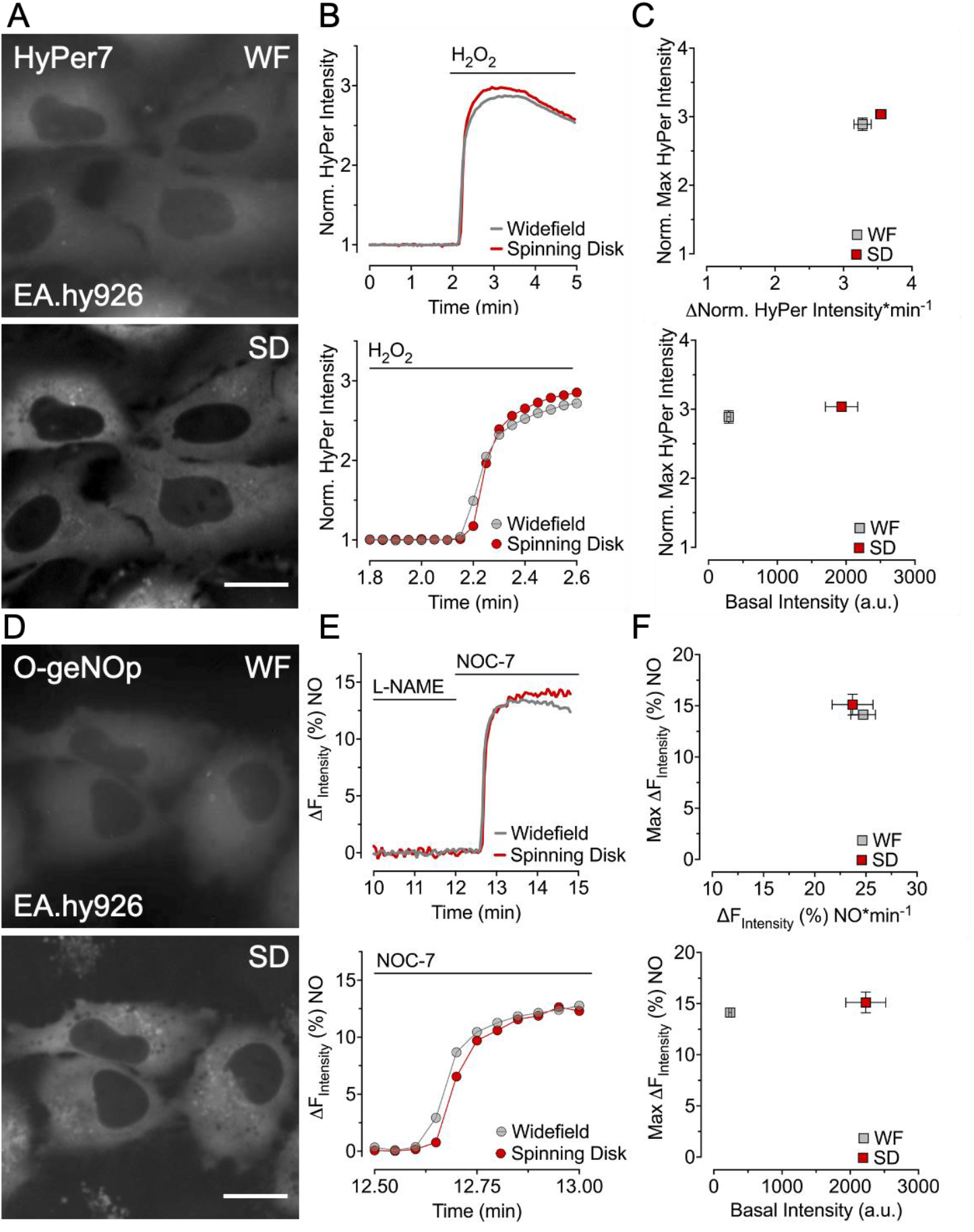
Comparison of HyPer7 and O-geNOp signals in the widefield and spinning disk mode. **(A)** Representative images of EA.hy926 cells stably expressing HyPer7 under different fluorescence microscopy modes widefield (WF) and spinning disk (SD). **(B)** Average curves show real-time H_2_O_2_ signals upon adding 25 μM of H_2_O_2_ in WF mode(grey line) or SD mode (red line). The lower panel is a close-up graphic of HyPer7 signals in a lower time frame. **(C)** In the XY scatter plot, the Y-axis shows the maxima of normalized HyPer7 fluorescence intensity, and the X-axis shows the rate of change in normalized HyPer7 intensity over time under different fluorescence microscopy modes widefield (WF) and spinning disk (SD). The lower panel shows an XY-scatter plot representing normalized HyPer7 fluorescence on the Y-axis and basal fluorescence intensity on the X-axis. Data are presented as mean ± SEM. (grey boxes: n=6/56, red boxes: n=6/56 WF and SD respectively) **(D)** Representative fluorescent images of O-geNOp in WF or SD mode. **(E)** Average curves show real-time NO signals upon adding 1 μM of NOC-7 using WF mode (grey line) or SD mode (red line). The lower panel shows the O-geNOp signals in a lower time frame. **(F)** XY scatter plot shows maximum NO responses in the Y-axis and rate of change in O-geNOp signals. In the lower panel in the XY-scatter plot, the Y-axis represents maximum NO responses, and X-axis represents basal fluorescence intensity. All data are presented as mean ±SEM (grey boxes n=4/37 and red boxes n=4/29 WF and SD, respectively). Scale bars represent 20 μm.

Statistical comparison of H_2_O_2_-induced HyPer7 signals measured with different microscopy settings showed that the maximum amplitude and the change in signal rate were comparable between SD and WF modes (Figure 3C, upper panel). Although the initial basal levels differed in both microscopy settings, the maximum responses were similar (Figure 3C, lower panel). Hence, both microscopy settings show comparable results in H_2_O_2_-induced HyPer7 activity measurements.

Next, we tested the O-geNOp signals in both imaging modes; administration of NOC-7 resulted in similar NO profiles and kinetics in both microscopy settings (Figure 3E). Additionally, we could not detect any differences between the SD and WF setup on NOC-7-induced O-geNOp maximum response and rate of increase in signals (Figure 3F, upper panel). Next, we assessed the relation between maximum O-geNOp responses and basal fluorescence intensity levels. Although basal fluorescence intensity was higher in SD mode, maximum responses of O-geNOp were similar in both modes (Figure 3F, lower panel). As a result, different microscopy settings showed similar results in NO-induced O-geNOp measurements.

Consequently, increasing the resolution of HyPer7 and O-geNOp biosensors with the spinning disk microscopy settings does not significantly affect the performance of the biosensors compared to widefield microscopy. These observations demonstrate that high spatial and temporal resolution can be achieved by camera-binning independently by employing a spinning disk microscope with camera binning setting 4×4.

### 2.4. Testing the role of incubation temperature on the performance of the biosensor

To understand the effect of temperature, we measured HyPer7 and O-geNOp signals in live cells at room temperature (RT) and 37°C, two conditions experimenters usually apply. First, we evaluated the effect of different ambient temperatures on the expression of biosensors. We observed that physiological temperature did not induce artifacts on the basal activity at the cellular level compared to RT either for Hyper7 or O-geNOp (Figures 4A and 4D). Activating HyPer7 by providing the cells with H_2_O_2_ at RT or 37°C did not significantly affect the performance of the biosensor (Figure 4B). Also, the maximum HyPer7 amplitude, the rate of signal increase, and the basal ratio remained unaffected under the two different ambient temperatures (Figure 4C). As a result, we conclude that HyPer7 activity remains agnostic to different temperatures in endothelial cells.

**Figure 4:**
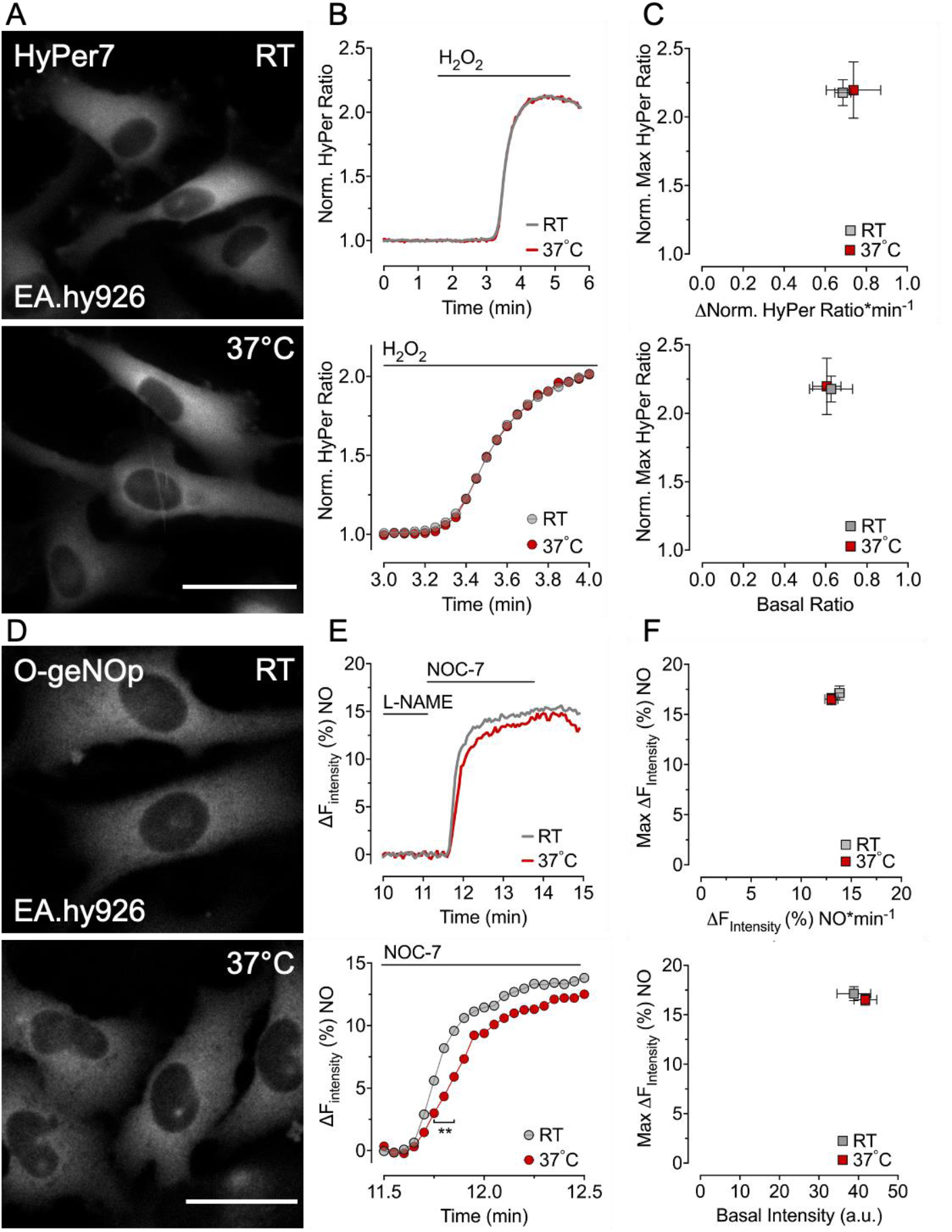
Measuring H_2_O_2_ and NO under different temperature conditions has no dramatic impact on responses of biosensors **(A)** Representative images of EA.hy926 cells stably expressing HyPer7 under room temperature (RT), and 37°C. **(B)** Average curves represent real-time H_2_O_2_ signals upon the addition of 25 μM of H_2_O_2_ in different temperatures (grey line RT, red line 37°C). The subjacent panel shows the close-up curve of HyPer signals upon the addition of H_2_O_2_. **(C)** In the XY scatter plot, Y-axis shows the maximum HyPer 7 responses, and X-axis represents the rate of change in the HyPer7 ratio over time. The lower scatter graphic plotted maximum HyPer7 responses against the basal HyPer7 ratio. All values are presented as mean ±SEM (grey boxes n=4/37 and red n=4/39 RT and 37°C, respectively). **(D)** Representative images of EA.hy926 cells stably expressing O-geNOp under room temperature (RT) and 37°C. **(E)** Average curves represent real-time NO signals upon the addition of 1 μM of NO and below close-up curves of O-geNOp signal in the lower time frame(Paired t-test *p-value* <0.01). The Grey curve represents measurements under RT conditions, and the red curve represents measurements under 37°C conditions. **(F)** In the XY plot, Y-axis represents the maximum responses of O-geNOp, and X-axis shows the rate of increase in O-geNOp signals. The lower XY-scatter plot shows maximum responses of O-geNOp in the Y-axis and basal fluorescence intensity of O-geNOp in the X-axis. All data presented as mean ±SEM (Grey n=4/40 and red n=4/37 represent RT and 37°C, respectively). Scale bars represent 20 μm.

Next, we employed the same experimental approach for O-geNOp responses. O-geNOp maximum response and NO profiles obtained at 37°C and RT were comparable (Figure 4E upper panel). However, the kinetics of O-geNOp signals showed a significantly slower increase in the initial on-kinetics at 37°C (Figure 4E, lower panel). Yet the maximum O-geNOp response, rate of increase, and basal fluorescence levels remained unaffected (Figure 4F). This observation might indicate a higher NO turnover and degradation at body temperature due to the increased enzymatic activity under physiological conditions. These data show that both biosensors suit multispectral imaging in endothelial cells at physiological or room temperature.

### 2.5. Multispectral imaging of HyPer7 and O-geNOp to visualize H_2_O_2_ and NO

Following the results from our previous optimization experiments, we decided to co-image both biosensors at RT, with a 4×4 camera binning on a conventional widefield microscope. Administration of NOC-7 instantly plateaued the O-geNOp signal while the respective H_2_O_2_ signal remained unaffected, indicating that exogenous NO does not acutely affect intracellular H_2_O_2_ (Figure S2). Subsequent provision of extracellular H_2_O_2_ was also ineffective in activating NO formation acutely, documented by O-geNOp signals (Figure S2). Changing the order of exogenous application of H_2_O_2_ first and then subsequently NOC-7 showed similar results (Figure S2), indicating that both reactants do not affect the intracellular generation of the other molecule.

Usually, intracellular levels of both reactive molecules, H_2_O_2_ and NO, are in a lower nanomolar range and are challenging to detect. Since pixel binning substantially improved H_2_O_2_ measurements, we co-imaged HyPer7 and O-geNOp with binning factors 4×4 versus 1×1. To test the endogenously generated NO and H_2_O_2_, we utilized histamine, a G-protein coupled receptor agonist, and Auranofin, a thioredoxin reductase inhibitor causing the accumulation of intracellular H_2_O_2_ [35]. Stimulating NO production with histamine increased the O-geNOp but not the HyPer7 signals (Figure 5A, left panel). Moreover, Auranofin only increased HyPer7 signals robustly but did not affect O-geNOp signals (Figure 5), documenting that under these optimized conditions, low levels of both reactive molecules can be accurately and dynamically detected (Figure 5A, left panel).

**Figure 5:**
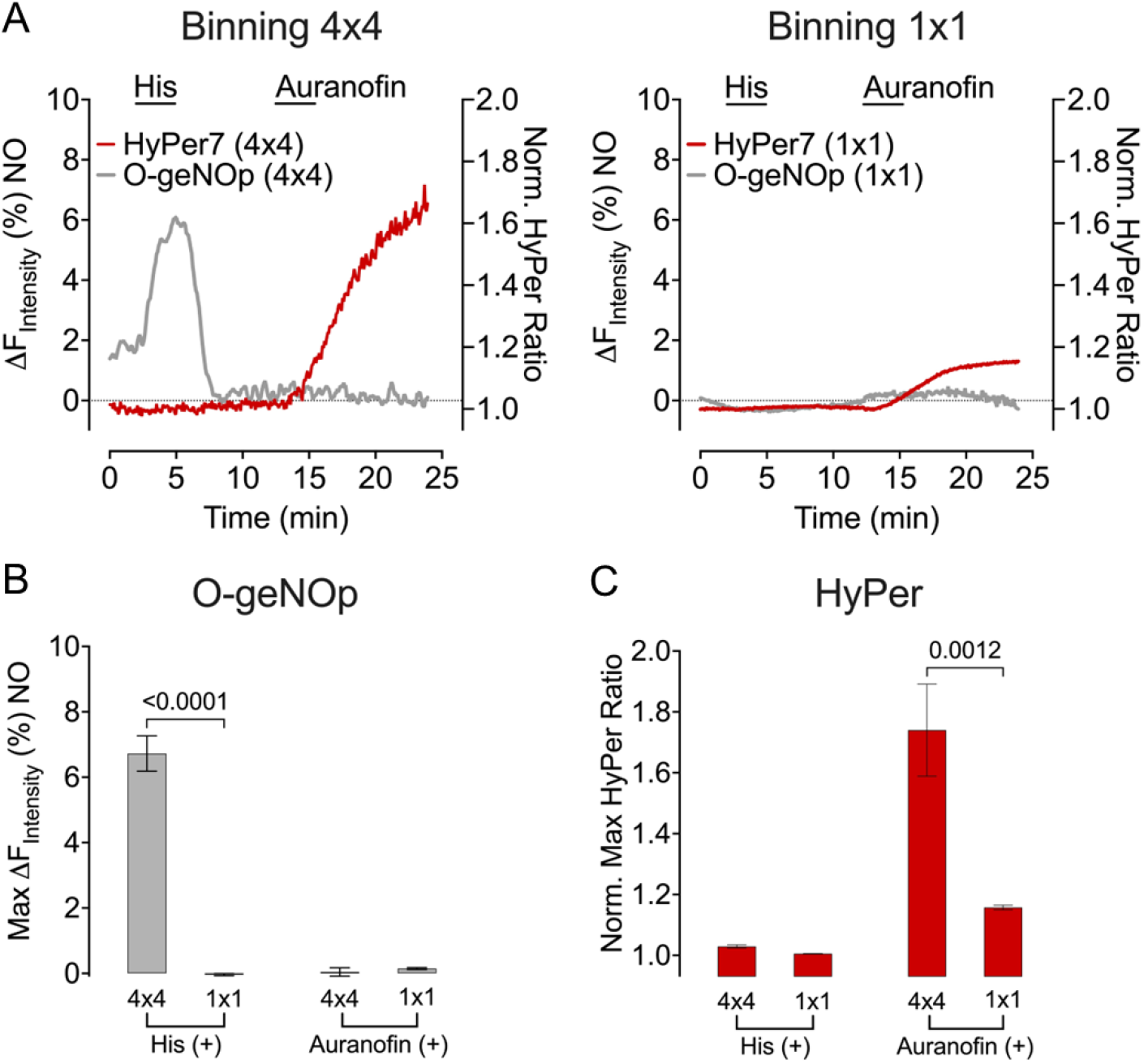
Simultaneous imaging of endogenously induced H_2_O_2_ and NO using different camera binning 1×1 and 4×4 **(A)** Average curves represent simultaneous measurements of HyPer7 (red lines) and O-geNOp (grey lines) in response to 30 μM histamine and 3 μM Auranofin, respectively. The left panel shows experiments performed with camera binning settings 4×4 and the right panel 1×1 **(B)** The bar plot shows the maximum responses of O-geNOp in response to histamine or Auranofin (4×4, n=3/38; 1×1, 3/17) **(C)** The bar plot shows the maximum responses of HyPer7 in response to histamine or Auranofin (4×4, n=3/38; 1×1, 3/17). All values are presented as mean SEM. Statistical significance was obtained using Student’s t-test, and p-values are indicated in bar plots.

The same experimental setup was applied with binning 1×1, resulting in significantly reduced H_2_O_2_ and completely diminished NO signals (Figure 5A, right panel, and Figure 5 B, C). Moreover, co-imaging O-geNOp with HyPer caused strong bleaching of the NO probe preventing the detection of even high levels of endogenous NO signals (Figure S3). These results demonstrate the importance of the binning factor when detecting low signals in a co-imaging mode using GEBs.

## 3. Discussion

In the present study, we pursued multispectral imaging techniques to simultaneously measure HyPer7 and O-geNOp signals in hardly transfectable endothelial cells. We have (i) generated double stable Ea.hy926 cells expressing two distinct biosensors, (ii) demonstrated that camera binning settings affect HyPer7 but not O-geNOp signals, (iii) proved that GEBs’ signal could be imaged equally efficiently on different microscope platforms, (iv) tested the role of ambient temperature on GEB functionality and (v) employed these optimized imaging settings to accurately co-image endogenous H_2_O_2_ and NO in endothelial cells.

The spectral distance between fluorescent biosensors HyPer7 and O-geNOp allows independent measurements of both analytes [36]. Hence, multispectral imaging is an excellent method for studying the direct interaction between H_2_O_2_ and NO in single endothelial cells. However, transient transfection methods are inefficient [33] in the endothelial cell line, EA.hy926 [37]. Hence, a transgenic methodology via viral transduction was used to generate cell lines expressing both HyPer7 and O-geNOp simultaneously (Figure 1). Functionality tests have shown that the probes are fully functional. Even though lentiviral vectors readily integrate into the host genome, their integration sites tend to be random [38]. Random integration of the open reading frame of a genetically encoded biosensor bears the risk of non-functional GEBs and may destroy endogenous gene expression patterns essential for cell function [39, p. 1], [40]. To limit these risks, instead of generating single double-stable clones of Ea.hy926 cells, we used polyclones to obtain a mixed cell population. One of the significant benefits of stable cell lines is that biosensor integration remains stable over long passages. Transient transfection of endothelial cells with conventional lipid-based transfection reagents is often toxic [41] to cells and may significantly alter their metabolism and signaling behavior [42]. Also, transient transfection does not allow long-term studies. Thus, future studies might employ gene editing approaches to selectively integrate the biosensors to so-called safe-harbor locations to ensure that the physiological state of cells is not altered.

Different biosensors have different dynamic ranges. HyPer’s dynamic range reaches up to a several-fold change in fluorescence upon total oxidation, while O-geNOp maximum response is limited to 15% (in this study). Therefore, optimal biosensor performance requires high SNR along with acceptable spatial resolution. A camera binning factor is a critical tool that can be tuned to obtain maximum signal dynamics. Investigating the effect of pixel binning on HyPer7 activity showed that low binning (higher resolution) negatively affected the HyPer7 dynamics (Figure 2). At high binning (lower resolution), the SNR of HyPer7 was increased because, at lower binning, the biosensor requires higher amounts of light to detect similar signals, which causes more phototoxicity at lower wavelengths (Figure S1). The phototoxicity effect was less pronounced for O-geNOp since its excitation is red-shifted and requires higher wavelengths (Figure 2). However, the phototoxicity became overt when O-geNOp recordings were performed simultaneously with HyPer7 at lower binning (high resolution) (Figure S3). Thus, we conclude that O-geNOp can be used at a lower binning factor when recorded in a single channel. For simultaneous measurements with HyPer7, lower resolution, and higher binning are required. Importantly, this observation might be transferable to other biosensors with similar spectral properties.

We next explored whether different imaging modes provide a compromise to obtain high-resolution and higher possible SNR and biosensors dynamic range. For this purpose, we investigated the effect of two modules, widefield and spinning disk. As expected, the spatial resolution of the images was higher with a spinning disk module than with the widefield module (Figure 3). Notably, the biosensor’s overall performance was comparable between widefield and spinning disk modes. Therefore, a high-resolution spinning disk module can be retained without losing signals of biosensor activity. High frame rates and fast camera acquisition allowed the spinning disk mode to obtain signals as sensitive as the widefield mode. Hence, the high resolution of spinning disk mode permits measuring local analyte concentrations by targeting the biosensor to subcellular compartments or analyzing samples of multiple cell layers without compromising signals.

While mammalian cells should ideally be kept at 37°C during GEB imaging, this may not be possible in all experimental setups. However, biosensor performance in different experimental setups may significantly vary [33], [34]. Thus, we tested the activity of Hyper7 and O-geNOp at room temperature and 37°C. Neither HyPer7 nor O-geNOp performance showed apparent alterations when tested at these temperatures. Only O-geNOp recordings showed a slightly retarded increase in the initial on-kinetic at 37°C (Figure 4), which might be attributable to a more vigorous intracellular enzymatic activity at physiological temperatures causing the faster degradation of the radical [43]. These results show that both biosensors can be employed at the given temperatures.

Next, we used these optimized parameters to simultaneously measure endogenous H_2_O_2_ and NO in human endothelial cells. For this purpose, we selected a conventional widefield microscope because these types of rigs are readily available to many laboratories and core facilities. Low binning (4×4) yielded sensitive HyPer and O-geNOp signals upon cell treatment with pharmacological tools compared to no binning (1×1) (Figure 5). Significantly, no binning did not affect O-geNOp functionality if measured in the OFP channel only (Figure 2G). Yet the capability of the NO probe to detect low levels of endogenous NO in endothelial cells completely diminished when measured in a co-imaging mode in a no-binning setting (Figure S3). These results may indicate that the green channel might strongly bleach the red-shifted biosensor due to exposure to high-energy wavelengths.

## 4. Conclusion

In conclusion, we generated a double stable cell line which allowed us to study the levels of H_2_O_2_ via HyPer7 and NO via O-geNOp simultaneously. We demonstrate that specific imaging parameters such as the camera binning, temperature, and imaging rig can dramatically alter the biosensor’s behavior. These changes may cause artificial signals, misinterpretation of data, or missing valuable signal information. Thus, we highly recommend testing critical parameters before establishing imaging protocols with the model systems of choice for different biosensors. The data and findings presented in this study may provide essential guidance for applying genetically encoded biosensors with similar features in live cells.

## 5. Materials and Methods

### 5.1. Molecular Cloning and Lentivirus production

Cytosolic targeted HyPer7 (HyPer7-NES), and O-geNOp (O-geNOp-NES) constructs were subcloned into a 3^rd^ generation lentivirus shuttle vector pLenti-MP2 (Addgene #36097), and HEK293T cells were used for lentivirus generation as described previously [23].

### 5.2. Cell Culture and Stable Cell Line Generation

Endothelial EA.hy926 cells were purchased from ATCC (CRL-2922, Manassas, VA, USA). Cells were maintained with Dulbecco’s minimal essential medium (DMEM)(Pan Bio-tech, Aidenbach, Germany) supplemented with 10% FBS (Pan Bio-tech, Aidenbach, Germany), 100 μg/mL Penicillin, 100 U/mL Streptomycin, 100 μg/mL Normocin (InvivoGen, San Diego, CA, USA), and 2% HAT ((Sodium Hypoxanthine (5 mM), Aminopterin (20 μM), and Thymidine (0.8 mM)) (ATCC, Manassas, VA, USA) in a humidified CO_2_ chamber (37°C, 5% CO_2_). EA.hy926 cells were seeded on a 6-well plate. When 50-60% confluency was achieved, cells were transduced with respective lentivirus HyPer7-NES and O-geNOp-NES using an antibiotic-free transduction medium containing 10% FBS and 10μg/ml Polybrene for 48-72h. Cells expressing both Hyper7-NES and O-geNOp-NES were selected using a fluorescence-activated cell sorter (FACS). The top 30% of the cells expressing both HyPer7-NES and O-geNOp-NES were sorted using a 488 nm laser (Filter: 530/40 nm) and 561 nm laser (Filter: 593/40 nm) on a BD Influx Cell Sorter. Sorted cells cultured as standard EA.hy926 culturing methods. Before the day of experiments, cells were seeded on a 6-well plate containing 30 mm coverslips (Glaswarenfabrik Karl Knecht Sondheim, Sondheim vor der Rhön, Germany).

### 5.3. Iron Loading Procedure

The cells were pre-treated with Iron (II) supplementation (300 μM FeSO4 and 500 μM ascorbate) for 15 min for full activation of the geNOp biosensor prior to experiments as described previously [44].

### 5.4. Buffers and Solutions

All chemicals were purchased from NeoFroxx unless otherwise stated. To maintain cells outside of the cell culture incubator, a cell storage buffer containing 2 mM CaCl_2_, 5 mM KCl, 138 mM NaCl, 1 mM MgCl_2_, 10 mM HEPES, 0.44 mM KH_2_PO_4_, 2.6 mM NaHCO_3_, 0.34 mM NaH2PO4, 10 mM D-Glucose, 0.1% MEM Vitamins (Pan-Biotech, Aidenbach, Germany), 0.2% essential amino acids (Pan-Biotech, Aidenbach, Germany), 100 μg/mL Penicillin (Pan-Biotech, Aidenbach, Germany), and 100 U/mL Streptomycin (Pan-Biotech, Aidenbach, Germany) was used. The pH was adjusted to 7.42 using 1 M NaOH. The cell storage buffer was sterile filtered with a 0.45 μm medium filter (Isolab, Germany). For live-cell imaging experiments, a HEPES-buffered solution was used consisting of 2 mM CaCl_2_, 5 mM KCl, 138 mM NaCl, 1 mM MgCl_2_, 10 mM HEPES, 10 mM D-Glucose, and pH was adjusted to 7.42 using 1 M NaOH. Histamine (Sigma-Aldrich, MO, USA) was prepared as 100 mM stock solution and diluted to 100 μM for imaging experiments using HEPES-buffered solution. Auranofin (Sigma-Aldrich, MO, USA) was prepared as 7.36 mM in DMSO for a stock solution and diluted to 3 μM for imaging experiments using a HEPES-buffered solution. NOC-7 (Sigma Aldrich, MO, USA) was prepared as 50 μM stock solution and diluted to 1-10 μM for imaging experiments using HEPES-buffered solution.

### 5.5. Imaging Experiments

#### 5.1.1. Widefield epifluorescence microscopy

Widefield epifluorescence microscopy experiments performed using Zeiss Axio Observer Z1.7 (Carl Zeiss AG, Oberkochen, Germany), Plan-Apochromat 20×/0.8 dry objective, Plan-Apochromat 40×/1.4 oil immersion objective, a monochrome CCD camera Axiocam 503, and a custom-made gravity-based perfusion system. The optical path for HyPer 7 starts with alternating excitation by 423/44 nm and 469/38 nm LED lights using a motorized filter wheel containing FT455(HyPer Low) and FT495 (HyPer High) beamsplitters(BS). Emissions were collected using the same emission filter (BP 525/50) in ratio imaging. For O-geNOp imaging, cells were excited with 555/30 nm LED light, and the filter combination is FT570 BS and 605/70 nm emission filter. The optical setup was the same, either mono-imaging or dual imaging of two biosensors. Data acquisition and control were set up using Zen Blue 3.1 Pro software (Carl Zeiss AG, Oberkochen, Germany). Provision and withdrawal of chemicals were conducted using an in-house gravity-based perfusion system connected to a perfusion chamber (NGFI, Graz, Austria)

#### 5.5.2. Spinning Disk Microscopy

Spinning disk microscopy experiments were performed on a Zeiss Axio Observer.Z1 equipped with both Yokogawa CSU-X1 (Tokyo, Japan) confocal scanner unit and Colibri 2 as light sources. This microscope is also equipped with LD A-plan 20x/0.3 dry objective and 2 different cameras, QuantEM:512SC (Teledyne Photometrics, AZ, USA) and AxioHrm, for confocal mode measurements and widefield measurements, respectively. HyPer7 signals were obtained by exciting cells with a 488 nm laser, and emission was collected at 509 nm in confocal mode. In widefield mode, cells were excited using 470/20 nm LED light, and the filter set contains FT495 BS, 525/25 nm emission filter. O-geNOp measurements were performed by exciting cells with a 558 nm laser, and emission was collected at 589 nm in confocal mode. In widefield mode, cells were excited using 550/12 nm LED light, and the filter set contains FT570 BS, 605/35 nm emission filter. Chemicals were provided or withdrawn using a gravity-based perfusion system connected to the perfusion chamber.

#### 5.5.3. Point Scanning Confocal Microscopy

Temperature experiments were performed on Zeiss AxioObserver equipped with LSM880 Confocal Laser Scan. To maintain and control ambient temperature microscope equipped with PeCon on-stage incubator(Erbach, Germany). For HyPer7 excitation, 405 nm and 488nm lasers were used, and for O-geNOp excitation, a 543 nm laser was used with a multibeamsplitter(MBS). The microscope is also equipped with 32 channel GaAsp detector to collect emissions. Cells were imaged using Plan-Apochromat 20x/0.8 M27 objective.

### 5.6. Data Analysis

For O-geNOp signals, background subtraction was performed using Microsoft Excel. Basal fluorescence intensities were analyzed employing a one-phase decay function in GraphPad Prism software to normalize O-geNOp signals to 100%. Raw fluorescence intensity is denoted as *F*, and normalized fluorescence intensity is denoted as *F_0_*. To obtain normalized signal curves formula used below:

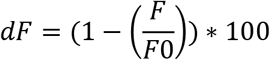

For HyPer signals, background subtraction was performed using Microsoft Excel. HyPer7 has two excitation and single emission termed HyPer low and HyPer high. The HyPer7 ratio was obtained by dividing HyPer high by HyPer low. Normalized HyPer ratios were obtained by normalizing HyPer ratios to their basal levels.

### 5.7. Statistical Analysis

All imaging data were analyzed using GraphPad Prism Software version 9 (GraphPad Software, San Diego, CA, USA). All experiments were performed at least three times and indicated as N/n where N represents the number of experiments and n represents the number of cells. All data represent with mean ± SEM unless otherwise stated. Statistical analysis of multiple groups was performed using one-way ANOVA with Tukey’s posttest (Comparison of all pairs of columns). P-values were indicated as numerical values. For the comparison of multiple groups, the p-value was not indicated if the p-value was higher than 0.05. In order to compare two experimental conditions unpaired Student’s t-test was performed. Statistical analysis of two groups was performed using unpaired Student’s t-test or Welch’s Student’s t-test, as indicated in the figures.

## Acknowledgments

Figure 1 was generated with BioRender.com (Agreement number: MI24VDYK0T).

## References

[1] R. Bretón-Romero and S. Lamas, ‘Hydrogen peroxide signaling in vascular endothelial cells’, Redox Biol, vol. 2, pp. 529–534, 2014, doi: 10.1016/j.redox.2014.02.005.

[2] A. K. Pandey et al., ‘Expression of CD70 Modulates Nitric Oxide and Redox Status in Endothelial Cells’, Arterioscler Thromb Vasc Biol, vol. 42, no. 9, pp. 1169–1185, Sep. 2022, doi: 10.1161/ATVBAHA.122.317866.

[3] S. S. Saeedi Saravi et al., ‘Differential endothelial signaling responses elicited by chemogenetic H2O2 synthesis’, Redox Biology, vol. 36, p. 101605, Sep. 2020, doi: 10.1016/j.redox.2020.101605.

[4] H. Sies, C. Berndt, and D. P. Jones, ‘Oxidative Stress’, Annual Review of Biochemistry, vol. 86, no. 1, pp. 715–748, 2017, doi: 10.1146/annurev-biochem-061516-045037.

[5] S. Steven et al., ‘Vascular Inflammation and Oxidative Stress: Major Triggers for Cardiovascular Disease’, Oxidative Medicine and Cellular Longevity, vol. 2019, p. e7092151, Jun. 2019, doi: 10.1155/2019/7092151.

[6] S. Rius-Pérez, I. Torres-Cuevas, I. Millán, Á. L. Ortega, and S. Pérez, ‘PGC-1*α*, Inflammation, and Oxidative Stress: An Integrative View in Metabolism’, Oxidative Medicine and Cellular Longevity, vol. 2020, p. e1452696, Mar. 2020, doi: 10.1155/2020/1452696.

[7] A. J. P. O. de Almeida et al., ‘Unveiling the Role of Inflammation and Oxidative Stress on Age-Related Cardiovascular Diseases’, Oxidative Medicine and Cellular Longevity, vol. 2020, p. e1954398, May 2020, doi: 10.1155/2020/1954398.

[8] H. Sies and D. P. Jones, ‘Reactive oxygen species (ROS) as pleiotropic physiological signalling agents’, Nat Rev Mol Cell Biol, vol. 21, no. 7, Art. no. 7, Jul. 2020, doi: 10.1038/s41580-020-0230-3.

[9] U. Förstermann and W. C. Sessa, ‘Nitric oxide synthases: regulation and function’, European Heart Journal, vol. 33, no. 7, pp. 829–837, Apr. 2012, doi: 10.1093/eurheartj/ehr304.

[10] J. Tejero, S. Shiva, and M. T. Gladwin, ‘Sources of Vascular Nitric Oxide and Reactive Oxygen Species and Their Regulation’, Physiological Reviews, vol. 99, no. 1, pp. 311–379, Jan. 2019, doi: 10.1152/physrev.00036.2017.

[11] A. R. Cyr, L. V. Huckaby, S. S. Shiva, and B. S. Zuckerbraun, ‘Nitric Oxide and Endothelial Dysfunction’, Crit Care Clin, vol. 36, no. 2, pp. 307–321, Apr. 2020, doi: 10.1016/j.ccc.2019.12.009.

[12] M. Gliozzi et al., ‘Modulation of Nitric Oxide Synthases by Oxidized LDLs: Role in Vascular Inflammation and Atherosclerosis Development’, International Journal of Molecular Sciences, vol. 20, no. 13, Art. no. 13, Jan. 2019, doi: 10.3390/ijms20133294.

[13] S. Thomas, S. Kotamraju, J. Zielonka, D. R. Harder, and B. Kalyanaraman, ‘Hydrogen Peroxide Induces Nitric Oxide and Proteosome Activity in Endothelial Cells: A Bell-Shaped Signaling Response’, Free Radic Biol Med, vol. 42, no. 7, pp. 1049–1061, Apr. 2007, doi: 10.1016/j.freeradbiomed.2007.01.005.

[14] G. C. Brown, ‘Reversible Binding and Inhibition of Catalase by Nitric Oxide’, European Journal of Biochemistry, vol. 232, no. 1, pp. 188–191, 1995, doi: 10.1111/j.1432-1033.1995.tb20798.x.

[15] H. Peng et al., ‘Nitric oxide inhibits endothelial cell apoptosis by inhibiting cysteine-dependent SOD1 monomerization’, FEBS Open Bio, vol. 12, no. 2, pp. 538–548, 2022, doi: 10.1002/2211-5463.13362.

[16] B. A. Maron and T. Michel, ‘Subcellular localization of oxidants and redox modulation of endothelial nitric oxide synthase’, Circ J, vol. 76, no. 11, pp. 2497–2512, 2012, doi: 10.1253/circj.cj-12-1207.

[17] U. Landmesser et al., ‘Oxidation of tetrahydrobiopterin leads to uncoupling of endothelial cell nitric oxide synthase in hypertension’, J Clin Invest, vol. 111, no. 8, pp. 1201–1209, Apr. 2003, doi: 10.1172/JCI14172.

[18] M. R. Depaoli et al., ‘Live cell imaging of signaling and metabolic activities’, Pharmacology & Therapeutics, vol. 202, pp. 98–119, Oct. 2019, doi: 10.1016/j.pharmthera.2019.06.003.

[19] E. Eroglu et al., ‘Genetic biosensors for imaging nitric oxide in single cells’, Free Radical Biology and Medicine, vol. 128, pp. 50–58, Nov. 2018, doi: 10.1016/j.freeradbiomed.2018.01.027.

[20] S. Okumoto, ‘Imaging approach for monitoring cellular metabolites and ions using genetically encoded biosensors’, Current Opinion in Biotechnology, vol. 21, no. 1, pp. 45–54, Feb. 2010, doi: 10.1016/j.copbio.2010.01.009.

[21] V. S. Ovechkina, S. M. Zakian, S. P. Medvedev, and K. R. Valetdinova, ‘Genetically Encoded Fluorescent Biosensors for Biomedical Applications’, Biomedicines, vol. 9, no. 11, p. 1528, Oct. 2021, doi: 10.3390/biomedicines9111528.

[22] A. E. Palmer, Y. Qin, J. G. Park, and J. E. McCombs, ‘Design and application of genetically encoded biosensors’, Trends in Biotechnology, vol. 29, no. 3, pp. 144–152, Mar. 2011, doi: 10.1016/j.tibtech.2010.12.004.

[23] M. Secilmis et al., ‘A Co-Culture-Based Multiparametric Imaging Technique to Dissect Local H2O2 Signals with Targeted HyPer7’, Biosensors (Basel), vol. 11, no. 9, p. 338, Sep. 2021, doi: 10.3390/bios11090338.

[24] V. V. Pak et al., ‘Ultrasensitive Genetically Encoded Indicator for Hydrogen Peroxide Identifies Roles for the Oxidant in Cell Migration and Mitochondrial Function’, Cell Metab, vol. 31, no. 3, pp. 642–653.e6, Mar. 2020, doi: 10.1016/j.cmet.2020.02.003.

[25] E. Eroglu et al., ‘Development of novel FP-based probes for live-cell imaging of nitric oxide dynamics’, Nat Commun, vol. 7, no. 1, Art. no. 1, Feb. 2016, doi: 10.1038/ncomms10623.

[26] E. Eroglu, H. Bischof, S. Charoensin, M. Waldeck-Weiermaier, W. F. Graier, and R. Malli, ‘Real-Time Imaging of Nitric Oxide Signals in Individual Cells Using geNOps’, Methods Mol Biol, vol. 1747, pp. 23–34, 2018, doi: 10.1007/978-1-4939-7695-9_3.

[27] E. Eroglu et al., ‘Application of Genetically Encoded Fluorescent Nitric Oxide (NO•) Probes, the geNOps, for Real-time Imaging of NO• Signals in Single Cells’, J Vis Exp, no. 121, p. 55486, Mar. 2017, doi: 10.3791/55486.

[28] E. Eroğlu, ‘Simultaneous Manipulation and Imaging of Chemogenetically Induced Hydrogen Peroxide in Hardly Transfectable Endothelial Cells’, Cumhuriyet Science Journal, vol. 43, no. 4, Art. no. 4, Dec. 2022, doi: 10.17776/csj.1114125.

[29] M. M. Frigault, J. Lacoste, J. L. Swift, and C. M. Brown, ‘Live-cell microscopy – tips and tools’, Journal of Cell Science, vol. 122, no. 6, pp. 753–767, Mar. 2009, doi: 10.1242/jcs.033837.

[30] C. M. Brown, ‘Fluorescence microscopy–avoiding the pitfalls’, Journal of Cell Science, vol. 120, no. 19, p. 3488, Oct. 2007, doi: 10.1242/jcs.022079.

[31] X. Ragàs, L. P. Cooper, J. H. White, S. Nonell, and C. Flors, ‘Quantification of Photosensitized Singlet Oxygen Production by a Fluorescent Protein’, ChemPhysChem, vol. 12, no. 1, pp. 161–165, 2011, doi: 10.1002/cphc.201000919.

[32] A. P. Wojtovich and T. H. Foster, ‘Optogenetic control of ROS production’, Redox Biology, vol. 2, pp. 368–376, Jan. 2014, doi: 10.1016/j.redox.2014.01.019.

[33] N. C. Shaner, P. A. Steinbach, and R. Y. Tsien, ‘A guide to choosing fluorescent proteins’, Nat Methods, vol. 2, no. 12, Art. no. 12, Dec. 2005, doi: 10.1038/nmeth819.

[34] E. Balleza, J. M. Kim, and P. Cluzel, ‘Systematic characterization of maturation time of fluorescent proteins in living cells’, Nat Methods, vol. 15, no. 1, Art. no. 1, Jan. 2018, doi: 10.1038/nmeth.4509.

[35] H. Hwangbo et al., ‘Auranofin Enhances Sulforaphane-Mediated Apoptosis in Hepatocellular Carcinoma Hep3B Cells through Inactivation of the PI3K/Akt Signaling Pathway’, Biomol Ther (Seoul), vol. 28, no. 5, pp. 443–455, Sep. 2020, doi: 10.4062/biomolther.2020.122.

[36] E. Eroglu, S. S. S. Saravi, A. Sorrentino, B. Steinhorn, and T. Michel, ‘Discordance between eNOS phosphorylation and activation revealed by multispectral imaging and chemogenetic methods’, Proceedings of the National Academy of Sciences, vol. 116, no. 40, pp. 20210–20217, Oct. 2019, doi: 10.1073/pnas.1910942116.

[37] C. J. Edgell, C. C. McDonald, and J. B. Graham, ‘Permanent cell line expressing human factor VIII-related antigen established by hybridization.’, Proc Natl Acad Sci U S A, vol. 80, no. 12, pp. 3734–3737, Jun. 1983.

[38] H. S. Kim et al., ‘CReVIS-Seq: A highly accurate and multiplexable method for genome-wide mapping of lentiviral integration sites’, Molecular Therapy - Methods & Clinical Development, vol. 20, pp. 792–800, Mar. 2021, doi: 10.1016/j.omtm.2020.10.012.

[39] S. Hacein-Bey-Abina et al., ‘LMO2-Associated Clonal T Cell Proliferation in Two Patients after Gene Therapy for SCID-X1’, Science, vol. 302, no. 5644, pp. 415–419, Oct. 2003, doi: 10.1126/science.1088547.

[40] M. Bokhoven et al., ‘Insertional Gene Activation by Lentiviral and Gammaretroviral Vectors’, Journal of Virology, vol. 83, no. 1, pp. 283–294, Jan. 2009, doi: 10.1128/JVI.01865-08.

[41] M. Bauer et al., ‘Toxic Effects of Lipid-Mediated Gene Transfer in Ventral Mesencephalic Explant Cultures’, Basic & Clinical Pharmacology & Toxicology, vol. 98, no. 4, pp. 395–400, 2006, doi: 10.1111/j.1742-7843.2006.pto_310.x.

[42] M. Danielli and R. A. Marinelli, ‘Lipid-based transfection reagents can interfere with cholesterol biosynthesis’, Analytical Biochemistry, vol. 495, pp. 1–2, Feb. 2016, doi: 10.1016/j.ab.2015.11.008.

[43] M. Kelm, ‘Nitric oxide metabolism and breakdown’, Biochimica et Biophysica Acta (BBA) - Bioenergetics, vol. 1411, no. 2, pp. 273–289, May 1999, doi: 10.1016/S0005-2728(99)00020-1.

[44] G. Sevimli et al., ‘Nitric oxide biosensor uncovers diminished ferrous iron-dependency of cultured cells adapted to physiological oxygen levels’, Redox Biology, vol. 53, p. 102319, Jul. 2022, doi: 10.1016/j.redox.2022.102319.

