## Supplementary for "Visualizing H_2_O_2_ and NO in endothelial cells: strategies and pitfalls"

### Supplementary Information

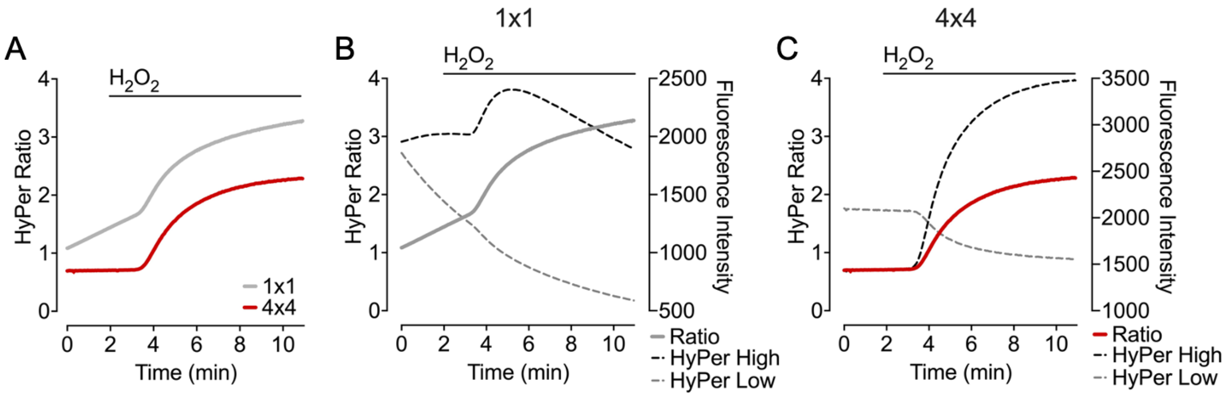


**Figure S1:** Effect of camera binning in HyPer7 raw signals. **(A)** Average curves represent HyPer7 ratios upon administration of 25 μM $H_2O_2$. The grey line represents the measurements using camera binning 1x1, and the red line represents measurements of 4x4 binning. **(B)** Average curves represent measurements of HyPer7 signals in 1x1 camera binning setup when cells were administered with 25 μM $H_2O_2$. The solid grey line shows the HyPer7 ratio signals, the dashed black line shows HyPer High (Ex/Em: 475/525) signals, and the dashed grey line shows HyPer Low (Ex/Em:430/525) signals. **(C)** Average curves represent measurements of HyPer7 signals in 4x4 camera binning setup when cells were administered with 25 μM $H_2O_2$. The solid grey line shows the HyPer7 ratio signals, the dashed black line shows HyPer High (Ex/Em: 475/525) signals, and the dashed grey line shows HyPer Low (Ex/Em:430/525) signals.

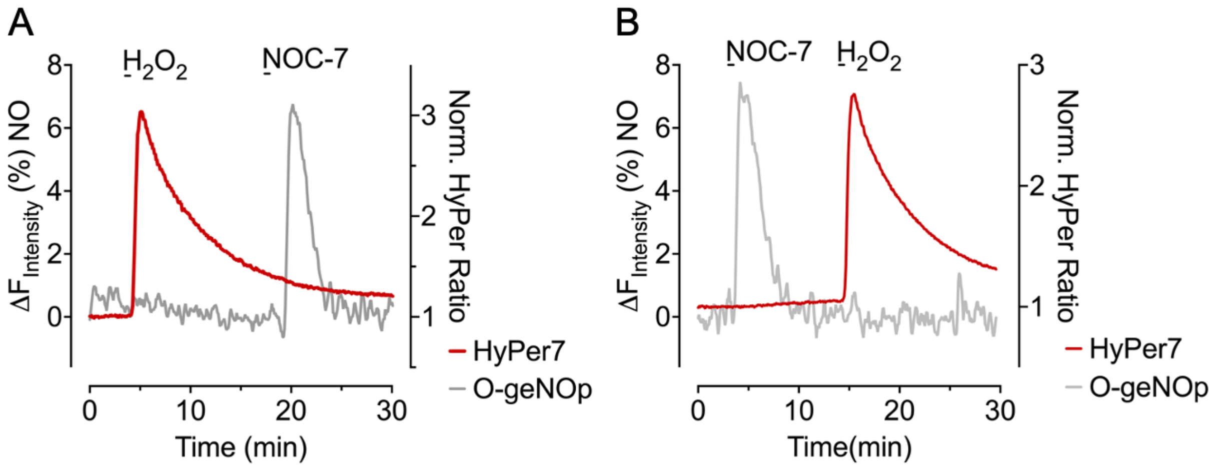


**Figure S2:** Simultaneous measurements of $H_2O_2$ and NO. **(A)** Average curve represents simultaneous measurements of HyPer7 (red line) and O-geNOp (grey line) by adding 1 µM NOC-7 and 25 µM $H_2O_2$, respectively. **(B)** Average curves represent simultaneous measurements of HyPer7 (red line) and O-geNOp (grey line) by adding 25 µM $H_2O_2$, 1 µM NOC-7, respectively.

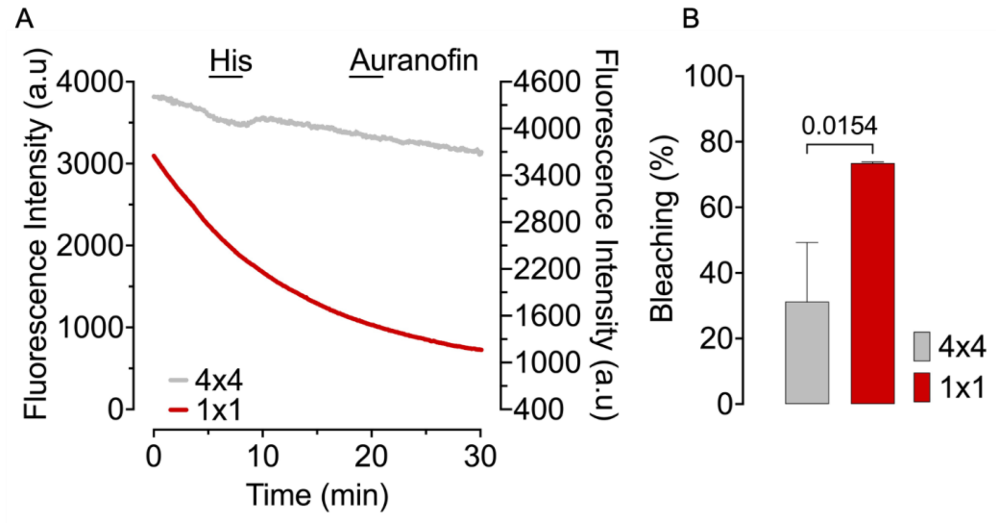


**Figure S3:** Co-imaging of O-geNOp with HyPer7 causes significant bleaching. **(A)** Representative curves show raw O-geNOp signals with different camera binning setups; 4x4 (grey line), 1x1(red line). **(B)** The bar plot shows the bleaching effect of O-geNOps signals in different camera binning setups. The grey bar showed bleaching in percentage when 4x4 binning factor was used. The red bar shows the bleaching of O-geNOp signals in 1x1 binning factor. All data were presented as mean ±SEM. Statistical analysis is performed using Student's t-test, and the p-value was indicated.
